# Poor biosafety and biosecurity practices and haphazard antibiotics usage in poultry farms in Nepal hindering antimicrobial stewardship

**DOI:** 10.1101/2023.04.17.536518

**Authors:** Ajit Poudel, Shreeya Sharma, Kavya Dhital, Shova Bhandari, Rajindra Napit, Dhiraj Puri, Dibesh B. Karmacharya

## Abstract

Poultry industry in Nepal has experienced remarkable growth in the last decade, but farm biosafety and biosecurity measures are often overlooked by farmers due to lack of knowledge or to save cost. As a result, farms often suffer from sporadic and regular outbreaks of many zoonotic diseases such as highly pathogenic avian influenza (HPAI), impacting production and creating public health challenges. Poor farm management practices, including overuse of antibiotics for prophylaxis and therapeutics, can complicate the spread of poultry diseases by creating and enhancing antimicrobial resistance (AMR) that is threating to both, poultry, and human health.

We assessed biosafety, biosecurity risks and AMR stewardship in sixteen poultry farms located in four districts (Ramechhap, Nuwakot, Sindhupalchowk, and Kavre) surrounding densely populated Kathmandu valley. Risk assessment and AMR stewardship evaluation questionnaire were administered to formulate biosafety and biosecurity compliance matrix (BBCM). Risk assessment checklist assessed facility operations, personnel and standard operating procedures, water supply, cleaning and maintenance, rodent/pest control and farm record keeping. Oral and cloacal samples from the poultry were collected, pooled, and screened for eight poultry pathogens using Polymerase Chain Reaction (PCR) tests.

Based on BBCM, we identified one of the farms in Sindhupalchowk (Farm 4) having the most (BBCM score= 67%) and a farm in Kavre (Farm 3) to having the least (BBCM score= 12%) biosafety and biosecurity compliance. Although most of the farms (61.6%) followed general poultry farming practices, only half had clean and well-maintained farms. Personal safety standard procedure compliance (BBCM score = 42.4%) and rodent control (BBCM score = 3.1%) were the biggest gaps. At least one of either bacterial or viral pathogen was detected in all farms. *Mycoplasma gallisepticum* was the most common disease detected in all but one farm, followed by *Mycoplasma synoviae*. Although more than half of the farmers considered AMR a threat, over 26% of them used antibiotics as a preventive measure and 81% did not consider withdrawal period for antibiotics prior to processing of their meat products. Additionally, antibiotics classified as Watch and Restrict by the WHO were frequently used by the farmers to treat bacterial infections in their farms. Lack of awareness and inadequate enforcement of regulations have exacerbated the risk of disease transmission in farms and compromised antimicrobial stewardship.

## Introduction

Nepal’s poultry industry has seen a significant and rapid growth in the last decade, contributing more than 4% to the national gross domestic product (GDP) [1,2]. Majority of the poultry products are supplied by numerous commercial farms (54% of total poultry production) scattered throughout the country. Backyard poultry also accounts for significant proportion of the total poultry production (46%); poultry meat and eggs are an easy source for protein and livelihood [1,3]. Rapidly expanding commercial poultry is reared in 64 out of 77 districts of Nepal and has an annual growth rate of over 18% [1,4]. According to the latest poultry census by the Nepal Central Bureau of Statistics [5], majority of chicken reared in Nepal are broilers (87%), with only small number of farms keeping layer chickens (11%). Almost half of the poultry production (46%) come from the central region of the country. Chitwan, Kathmandu, and Kaski districts account for more than 85% of total meat and eggs production.

In Nepal, in spite of burgeoning poultry industry, proper biosecurity measures are often overlooked [6]. Backyard poultry farmers often feel the burden of maintaining biosecurity due to lack of knowledge and perceived additional cost [7]. Biosafety encompasses measures to prevent transmission of infectious diseases, and biosecurity measures are meant to prevent introduction and spread of pathogens in farms. By implementing a proper biological containment (and exclusion) along with traffic control, segregation, and sanitation- an effective biosafety and biosecurity can be maintained [8]. Keeping healthy flocks not only guarantees financial security for the farmers, it can also prevent outbreaks of zoonotic diseases such as HPAI [2,9]. With increased commercial poultry production, maintaining biosecurity and biosafety measures in farms have been challenging [6]. Lack of government initiatives (and efforts), both in developing and developed countries alike, to raise awareness and implement regulations on biosecurity and biosafety have also grossly undermined proper safe farming practices [10]. Although Veterinary Standards and Drug Administration Office (VSDAO) in Nepal has developed a manual for proper poultry management including biosecurity guidelines, it isn’t properly enforced and is often overlooked by farmers [11]. Poultry production that are primarily focused on profitability, with compromised biosafety and biosecurity practices for cost saving, will eventually face production loss and increased health risks to both birds as well as humans (and other animals) [8].

Despite increasing occurrence of disease outbreaks such as Avian Influenza (AI) in poultry farms, many farmers in Nepal are unaware of the importance of biosecurity measures and their practices [7]. This lack of awareness, along with inadequate enforcement of biosecurity regulations, is exacerbating the risk of AI transmission in the country [12]. Poor farm management practices such as the use of unprocessed poultry waste as animal feed and the overcrowding of birds due to poorly designed farm contribute to the spread of AI [13].

Disease outbreaks in poultry farms due to bacterial pathogens such as *Mycoplasma gallisepticum* (Mg), *Mycoplasma synoviae* (Ms), *Escherichia Coli* (E. coli) and salmonella also occur due to lapses in biosafety and biosecurity. Rampant and haphazard use of antibiotics in poultry farms in Nepal is a leading cause of antimicrobial resistance (AMR) and a looming threat to human health [14]. There is a need to reduce and promote responsible use of antibiotics in the poultry industry in Nepal [15,16]. Antimicrobial stewardship, which is a coordinated program that promotes the appropriate use of antimicrobials (including antibiotics), reduces microbial resistance, and decreases the spread of infections caused by multidrug-resistant organisms, has become one of the important aspects of a comprehensive biosecurity and biosafety practices [7].

Kathmandu is a densely-populated metropolitan capital city of Nepal with a population of over 2 million people [17]. Any emerging, re-emerging and diseases of human health concern originating from animal production sites, such as poultry farms, can rapidly spread in a city like Kathmandu. Since disease outbreak, transmission and spread dynamics are directly linked to biosafety and biosecurity status of farms, it is vital to understand the current status of the farms located close to the city. We conducted a comprehensive risk assessment and status evaluation of biosafety, biosecurity and AMR stewardship in sixteen poultry farms located in four districts with high poultry production (Ramechhap, Nuwakot, Sindhupalchowk, and Kavre) surrounding the Kathmandu valley.

## Methodology

### Study Site and Data collection

Four farms (small and medium sized; poultry <2000 per farm) in each of the four districts (Kavre, Sindhupalchowk, Ramechhap and Nuwakot) were selected for the study (Figure 1). Risk assessment checklist (Supplementary Table 1) and AMR stewardship questionnaire (Supplementary Table 2) were used to collect data. Risk assessment checklist assessed facility operations, personnel and standard operating procedures, water supply, cleaning and maintenance, rodent/pest control and farm record keeping.

**Figure 1:**
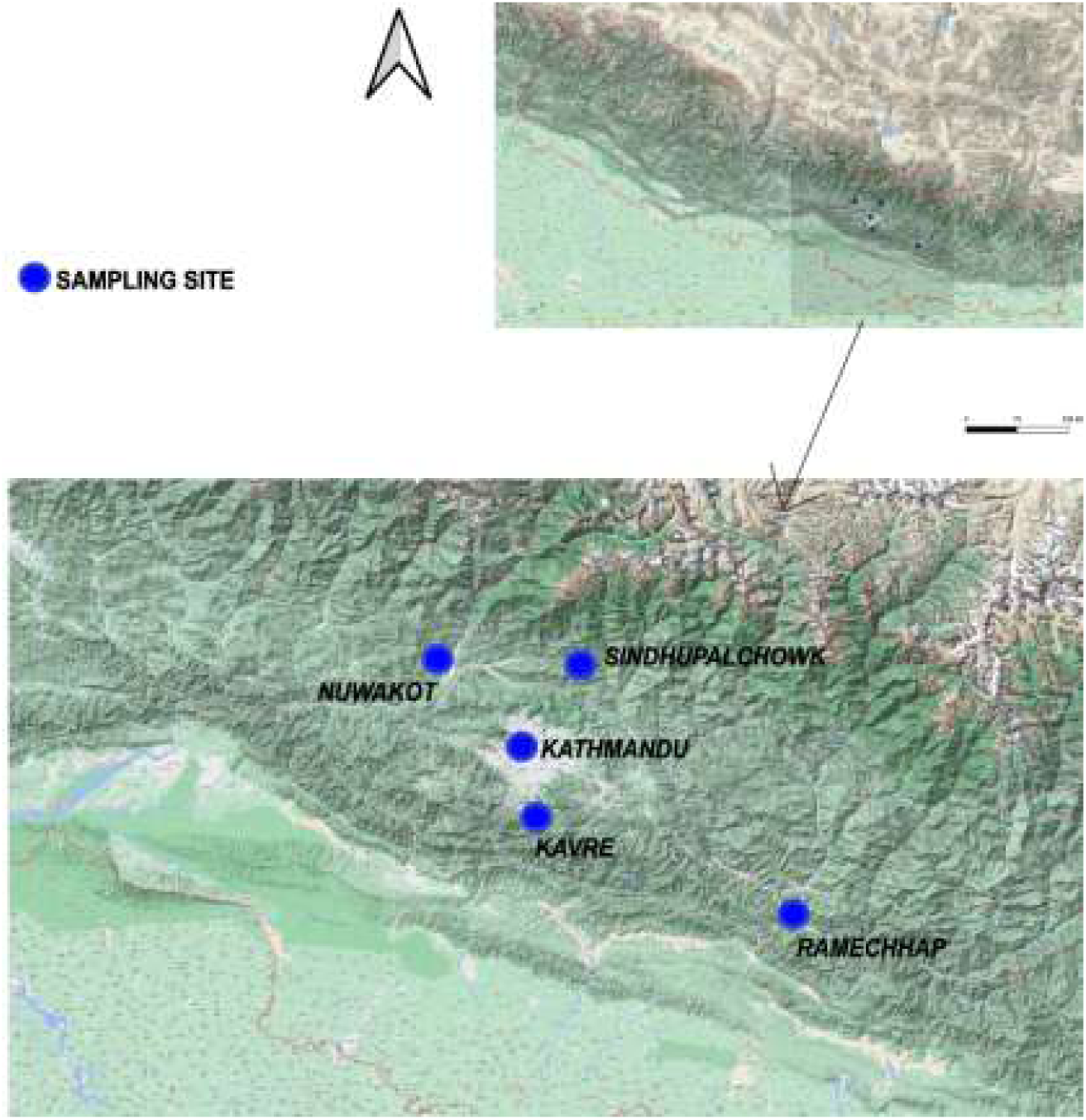
Selected poultry farms in the districts (Kavre, Sindhupalchowk, Ramechhap and Nuwakot) surrounding Kathmandu valley. The top map shows location of the districts in Nepal (shown in red border) and the bottom map shows their location in respect with Kathmandu Valley (shown just for reference, not a sampling site). The map was generated using QGIS software, Version 3.30.0 [18].

### Biological Sample Collection

Oropharyngeal (n=1) and cloacal (n=1) swabs from each chicken were collected from selected farms (n= 16 farms; 30 birds per farm) and stored in viral transport media (VTM). All samples were stored in ice boxes (2- 8^0^ C) during sample collection and transported in liquid nitrogen container (-196^0^ C) to BIOVAC’s lab in Kathmandu. The samples were then pooled (n=10) from each farm (n= 6 pooled samples; pooled oropharyngeal swabs=3 and pooled cloacal swabs=3). These pooled samples were screened for eight poultry pathogens- Newcastle disease virus (NDV), Influenza A Virus (IAV), Infectious Bronchitis Virus (IBV), Infectious Bursal Disease (IBD), *Mycoplasma synoviae (Ms)*, *Mycoplasma gallisepticum (Mg)*, Marek’s Disease Virus-1 (MDV1) and Marek’s Disease Virus-2 (MDV2) using PCR.

### Nucleic Acid Extraction and PCR

The nucleic acid (DNA/RNA) from pooled samples were extracted using automated nucleic acid extractor (abGenix™ AITbiotech, Singapore) following manufacturer’s instructions. The pooled swab samples were stored at -20°C. PCR for detection of the eight pathogens were performed using SuperScript™ III Platinum™ One-Step qRT-PCR Kit w/ROX (Invitrogen, Catalog number 11745500). The primers for NDV, IBV, IBDV, Mg, Ms, MDV1 and MDV2 were designed using NCBI PrimerBlast® (Table 1).

**Table 1:**
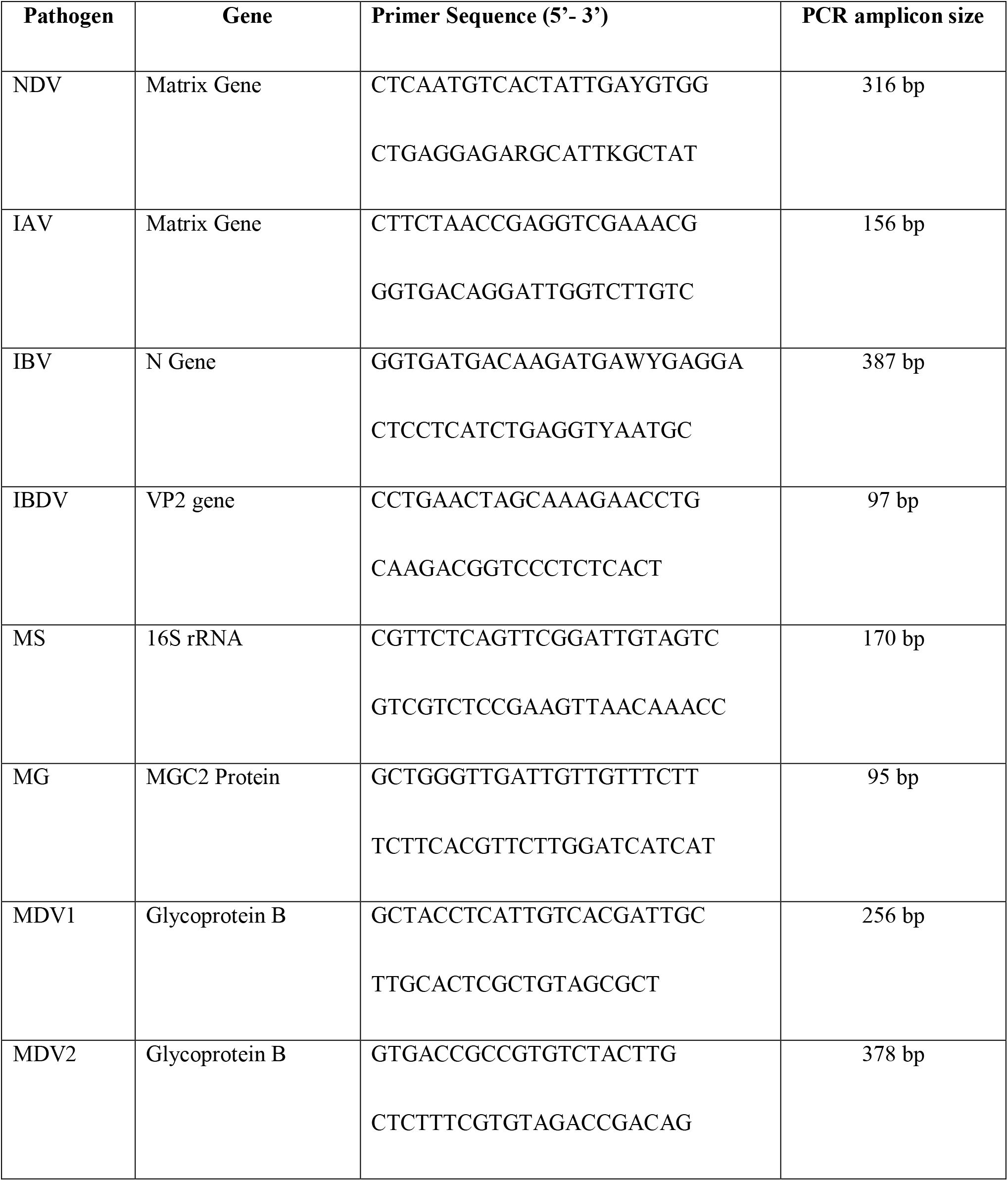
PCR primers for each of the pathogen designed using PrimerBlast®.

For IAV, the primers IAV ISO_F and IAV ISO_R were used (Table 1) [19].

### PCR Condition

PCR for each pathogen test was done in a 25ul reaction containing 4 μL of extracted RNA (for NDV, IAV, IBD, IBV) and 4 μL of extracted DNA (for MG, MS, MDV1 and MDV2), 1 μL each of respective 10pm forward and reverse primers, 12.5 μL of 2X Mastermix with ROX, 0.2 μL of SuperScript™ III Platinum™ enzyme and 6.3 μL of Nuclease free water. All eight PCR were carried out with enzyme activation at 45°C for 15 minutes followed by one cycle of initial denaturation at 95°C for 5 minutes. PCR for RNA viruses consisted of 45 cycles of denaturation at 95°C for 30 seconds, annealing at 59°C for 30 seconds and extension at 72°C for 20 seconds. PCR for MS, MG, MDV1 and MDV2 consisted of 10 cycles of denaturation at 95°C for 30 seconds, annealing at 63°C for 40 seconds and extension at 72°C for 20 seconds followed by 35 cycles of denaturation at 95°C for 30 seconds, annealing at 60°C for 40 seconds and extension at 72°C for 20 seconds. The final extension for all eight PCR were carried out at 72°C for 2 minutes. All the amplified PCR products were visualized using 1.5% Agarose Gel (Supplementary Figures).

### Biosafety and Biosecurity Risk Assessment

#### Biosafety and Biosecurity Compliance Matrix (BBCM) Score

We created a Biosafety and Biosecurity Compliance Matrix (BBCM) score based on risk assessment checklist which included criteria such as facility operations, personal and standard operating procedures, water supply, cleaning and maintenance, rodent/pest control and farm record keeping (Supplementary Table 1). General practices relate to infrastructure of or within the farm that aids in its biosecurity. Personnel standards and procedures include hygiene practices, use of disinfectants and precautionary measures taken to minimize pathogen contamination in farms. Water supply assessed whether the water given to poultry is adequately disinfected and clean. We also investigated preventive measures taken by farms to control rodents. Poultry rearing area and equipment cleanliness practices implemented by the farm personnel were also evaluated under cleaning and maintenance criteria. And finally, under record keeping criteria, we assessed practices of keeping records of daily activities, material usage/ consumption and any breaches detected in the farm.

For each selected criteria complied, one point was given for every activity implemented and a BBCM score was tallied for each category in every farm. A final farm BBCM score was then calculated and converted into percentage. We categorized farms that had >90% BBCM score as high, 60-89% as medium and <60% as low. Scores received by each farm are shown in Table ***2*** and visualized in Figure 2.

**Figure 2:**
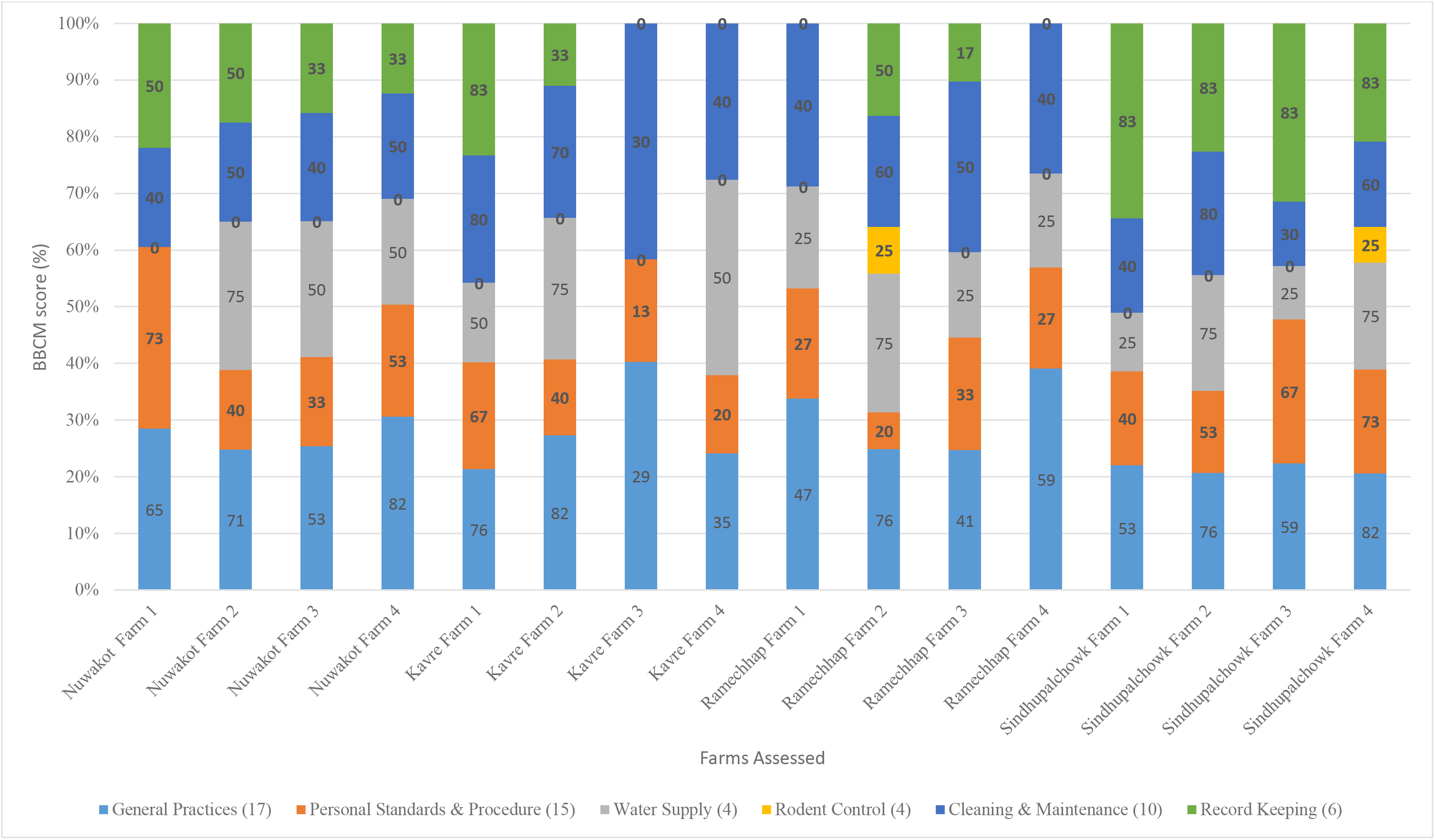
Graphical representation of BBCM scores received by all 16 farms for various biosafety & biosecurity parameters. Scores (percentage) received by all farms assessed in this study using BBCM score. Numbers in parenthesis in the legend refers to total number of activities assessed within each criterion.

**Table 2:**
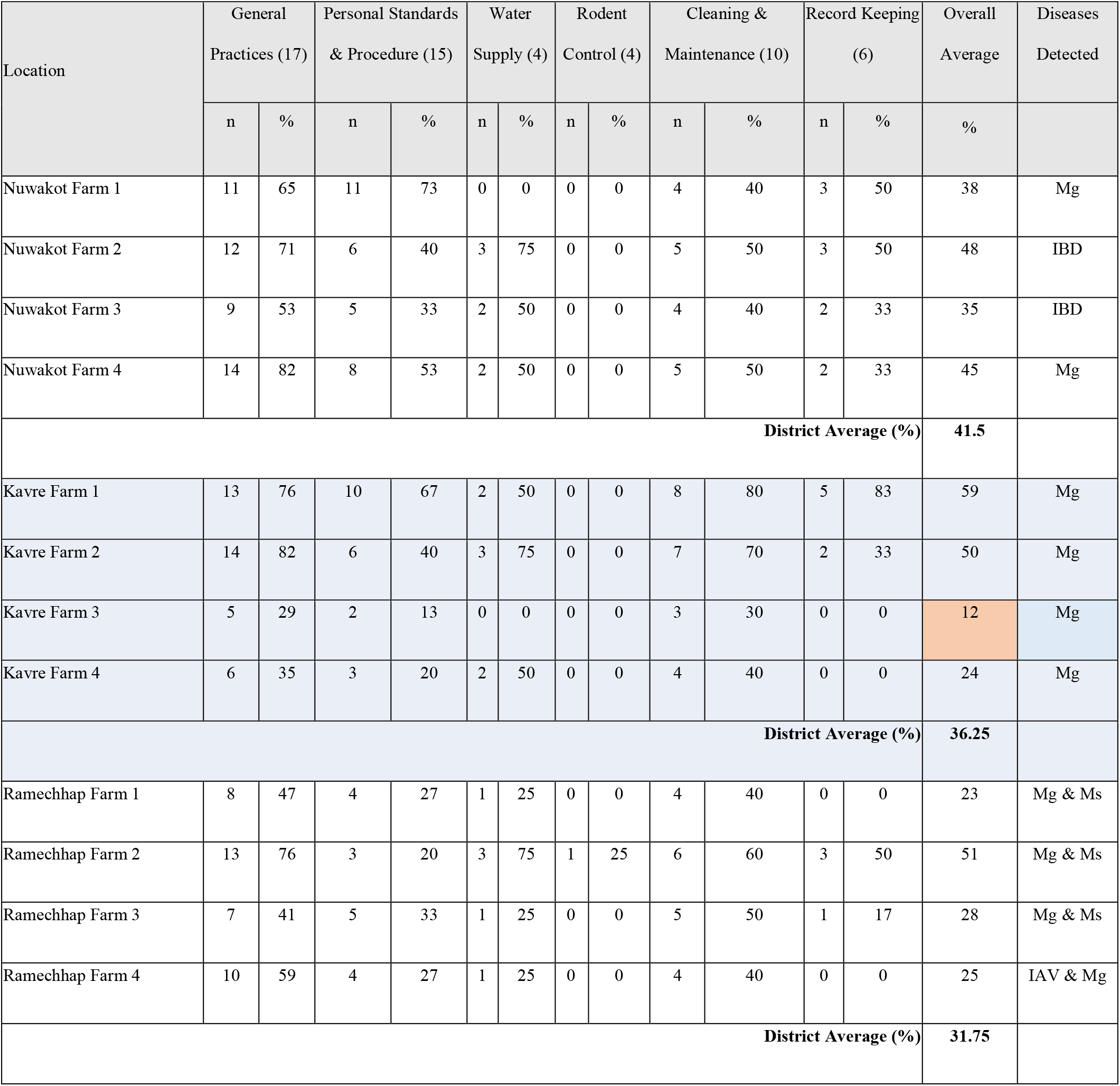
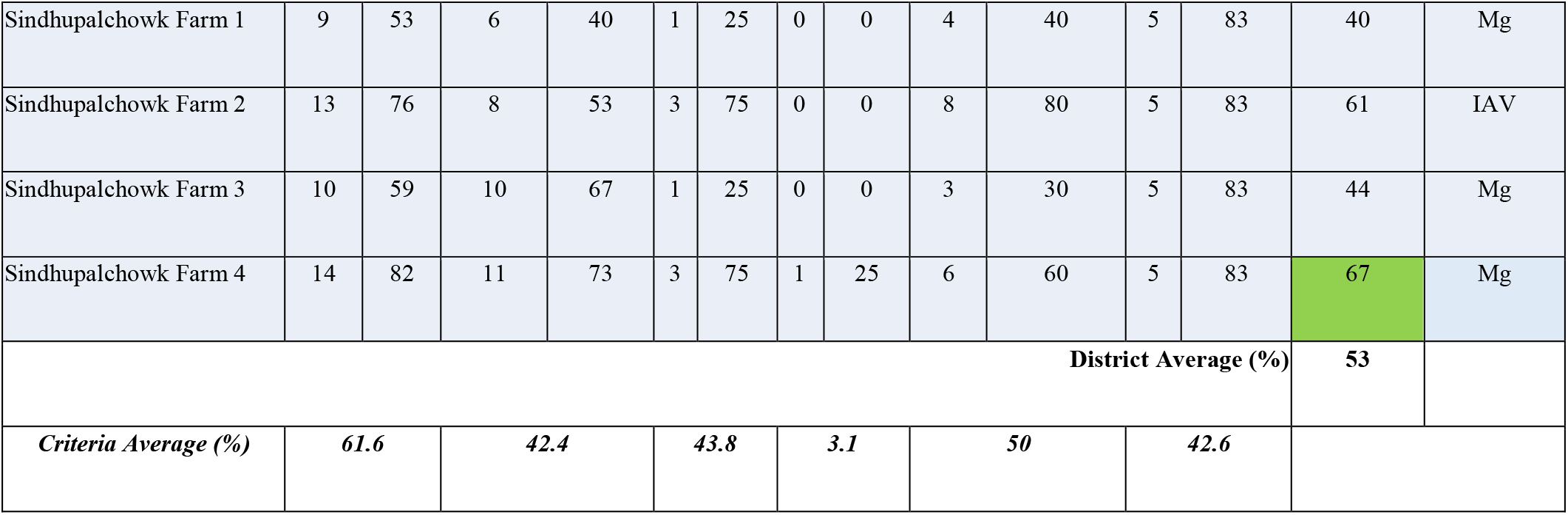
Total number and percentage of BBCM criteria met by the farms as per our assessment checklist. Overall average BBCM score was calculated by dividing the sum of all BBCM fulfilled criteria (n) and calculating the percentage. The highest BBCM score (Green cell- Sindhupalchowk Farm 4) was 67%, and the lowest BBCM score was 12% (Orange cell- Kavre Farm 3). The total number of activities assessed in each criterion are listed within parenthesis. Details of the activities assessed are shown in Supplementary Table 1. Mg – Mycoplasma gallisepticum, Ms – Mycoplasma synoviae, IBD – Infectious Bursal Disease, IAV – Influenza A Virus

## Results

### Biosafety and Biosecurity Risk Assessment

Analysis of overall Biosafety and Biosecurity of all farms surveyed showed low compliance (average BBCM score= <41%). The highest BBCM rating was scored by a farm in Sindhupalchowk (Farm 4, BBCM= 67%) and the lowest was by a farm in Kavre (Farm 3, BBCM= 12%). At district level, Sindhupalchowk had the most Biosafety and Biosecurity compliance (BBCM= 53%) whereas Ramechhap had the least (BBCM= 32%). Of all the assessed criteria, rodent control was the most neglected (BBCM= 3.1%). Only two farms (Ramechhap Farm 2 and Sindhupalchowk Farm 4) had implemented one out of four record keeping practices. General poultry farming practice was the most biosafety and biosecurity compliant criteria fulfilled by all farms (BBCM= 61.6%) (Table 2).

General practices, personal standards and procedure, and cleaning and maintenance were the only categories implemented in all sixteen farms (Table 2, Figure 2). Only two farms (Ramechhap 2 and Sindhupalchowk 4) implemented rodent control, these were also the only farms that implement some activities of all six criteria. Water supply was implemented in all but two farms (Nuwakot 1 and Kavre 3). Similarly, record keeping was implemented in all but four farms (Kavre 3, Kavre 4, Ramechhap 1, and Ramechhap 4).

Across all 16 farms, general practices received the highest criteria average (61.6%), followed by cleaning and maintenance (50%), water supply (43.8%), record keeping (42.6%), personal standards and procedure (42.4%), and rodent control (3.1%).

At least one of either screened bacterial or viral pathogen was detected in all farms. Mg was the most common disease detected, in all but one farm, followed by Ms. Farms in Ramechhap district were most affected, with Mg detected in all four farms, Ms in three, and IAV in one. Two different poultry diseases (IAV and Mg) were detected in Sindhupalchowk (Farm 2). IBD (Farm 2 and 3) and Mg (Farm 1 and 4) were detected in Nuwakot district (Table 2). Results of PCR tests detecting the diseases (gel images) are shown in Supplementary Figures.

### AMR Stewardship

A little over half (52%) of the surveyed farmers considered AMR as a real threat (Table 3). Almost all farmers (93%) had not attended any programs or campaigns related to AMR. Very few farmers (12%) received training on AMR stewardship; these farmers received training/information on AMR from local animal health centres (21%), veterinarians (29%), vet technician (17%) or from vet suppliers (21%). Majority of the farmers (81%) did not consider implementing “withdrawal period” for antibiotics use prior to selling their meat products. Most of the famers trusted veterinarians (40%) and vet technician (14%) on receiving consultation on antibiotics use. Farmers used various antibiotics for prophylaxis (26%) and therapeutics (76%) needs.

**Table 3:** Knowledge and perception of AMR among poultry farmers. . Responses (in percentage) obtained from farmers on AMR-related questions.

| AMR Questions |  | Percentage |
| --- | --- | --- |
| Is AMR a threat? | Yes | 52 |
|  | No | 48 |
| Where did you get AMR knowledge from? | Local Animal Health Center | 21 |
|  | Veterinarian | 29 |
|  | Vet Technician | 17 |
|  | Vet Suppliers & Shops | 21 |
|  | Self | 12 |
| Who do you most trust for consultation? | Veterinarians | 40 |
|  | Vet Technician | 14 |
|  | Self | 2 |
|  | Local Animal Health Center | 26 |
|  | Shops & Sales Team | 14 |
|  | Local Animal Health Center | 2 |
| Do you consider withdrawal period after antibiotics use? | Yes | 12 |
|  | No | 81 |
|  | Don't Know | 7 |
| Withdrawal period prior to culling? | Yes | 14 |
|  | No | 79 |
|  | Don't Know | 7 |
| Any programs/campaigns<br>attended on AMR? | Yes | 5 |
|  | No | 93 |
|  | Don't Know | 2 |
| Antibiotics used for<br>prophylaxis? | Yes | 26 |
|  | No | 74 |
| Antibiotics used for<br>therapeutics? | Yes | 76 |
|  | No | 24 |

The farmers were also inquired about various types and proportion of antibiotics they used (Figure 3). Tetracycline was the most used (36%) antibiotics, followed by Colistin (14%), Quinolone (12%), Aminoglycoside (12%) and Macrolide (9%). Penicillin (3%) and Beta-lactam (2%) were the least used antibiotics.

**Figure 3:**
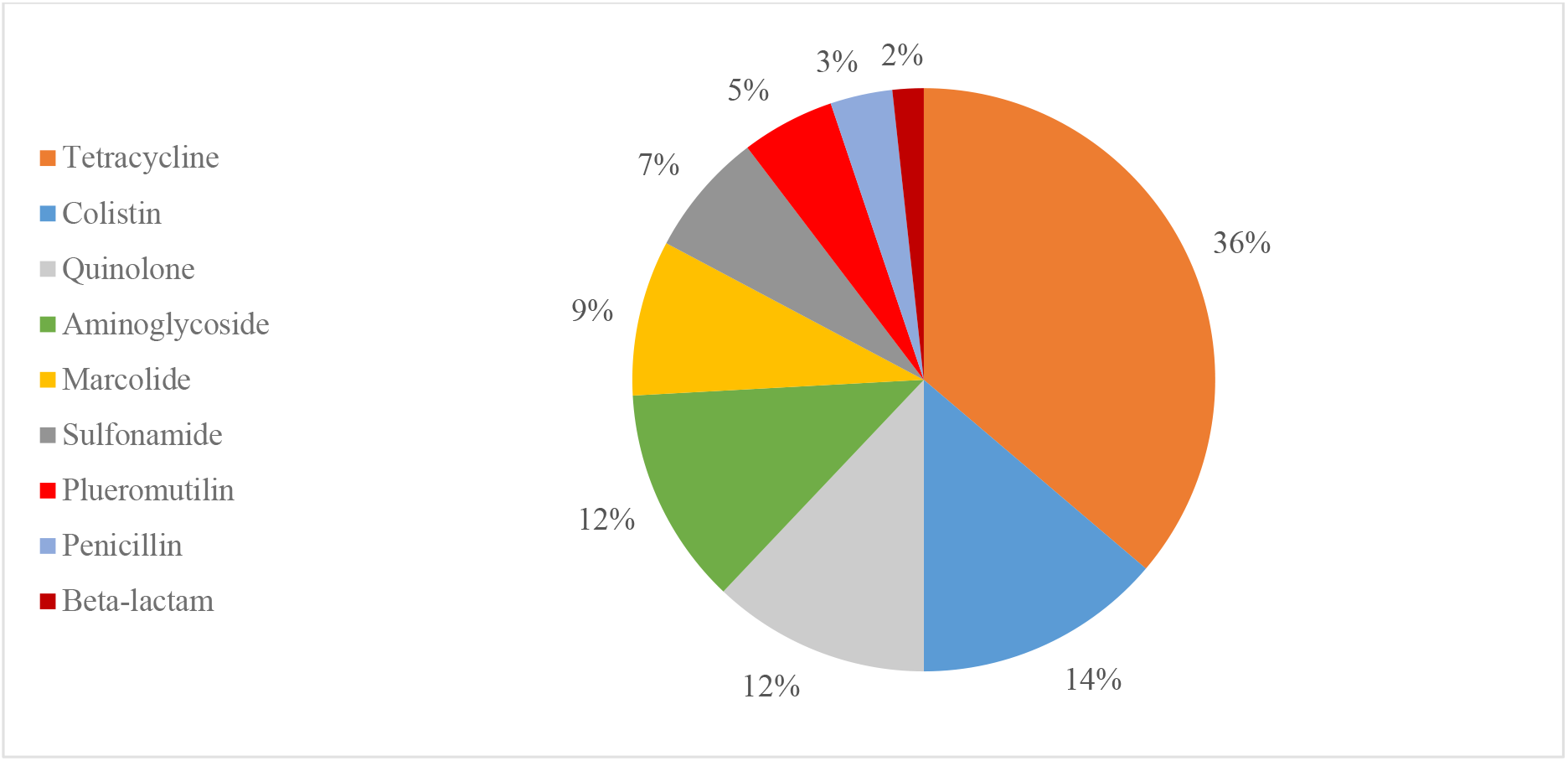
Various types of antibiotics used (in %) in the surveyed poultry farms.

## Discussion

Poor biosafety and biosecurity measures on poultry farms can lead to introduction and spread of bacterial and viral diseases. Regular monitoring and testing, an important component of implementing strict biosecurity and biosafety measures, are essential for detecting and containing infections in farms. Poultry farms with compromised biosafety and biosecurity compliance often suffer from various prevalent bacterial and viral infections such as Mycoplasma, IBD and IAV- affecting overall poultry health and lowering production.

Mg and Ms are bacterial diseases which spread through direct contact, respiratory secretions, or contaminated materials [20]. IBD, on the other hand, is a viral disease that can cause damage to the immune system of young chickens, often leading to mortality [21]. It is transmitted through contact with contaminated sources, such as faeces. IAV in poultry is primarily transmitted through direct contact with infected birds or through contact with contaminated surfaces, feed, or water. Wild birds, particularly waterfowl, are the natural reservoirs for the virus and are thought to be one of the major sources of avian influenza outbreaks [22,23].

Introduction of infected birds, equipment, or materials in farms can result in the spread of diseases [24,25]. Movement of people and animals on and off the farm can also contribute to the spread of disease [26]. Birds that are housed in crowded or unsanitary conditions are susceptible to diseases; maintaining proper biosecurity and biosafety measures in such farms are often challenging [27]. Implementation of these measures were severely lacking in the farms that we surveyed in our study. Only one farm scored medium biosafety and biosecurity compliance rating (Sindhupalchowk Farm 4, BBCM= 67%). None of the farms received a high score of >90%, and not surprisingly, diseases were detected in all farms (Table 2). Moreover, out of all sixteen farms, only two (Ramechhap 2 and Sindhupalchowk 4) had evidence of conducting activities pertaining all six different categories of the assessment. Rest of the farms had activities/practices missing from an entire category.

Biosecurity measures, such as controlling the movement of people and animals onto the farm, implementing disinfection procedures, and providing protective clothing, can significant improve poultry production by reducing disease outbreaks and spread [28,29]. We detected Mg and Ms in most of the farms, these bacterial infections have low mortality but associated morbidity directly affects egg and meat production [30]. Overall, the farms in Ramechhap district scored poorly in BBCM, something the district animal health authority needs to be aware of and make effort to improve.

Although majority of the farms focused on general practices, cleaning and maintenance, and personal standards and procedures, except for two farms (Ramechhap 2 and Sindhupalchowk 4) most had completely ignored rodent control (Table 2). Rodents are a major pest and a disease vector of poultry farms [31]. Rodents are also known to cause damage to farm infrastructure and feed, kill young chicks and eat eggs [32].

Poultry farms often resort to using medications to treat and salvage their flock only after disease outbreaks. We detected viral pathogens (IBD and IAV) and bacterial infections (Mg and Ms) in all but one farm (out of 16). Antibiotics was rampantly used, both as prophylaxis and therapeutics, by the farmers. Antibiotics such as Tetracycline, Colistin, Quinolone, Aminoglycoside, and Macrolide were the most used antibiotics by the farmers- consistent with national trend [33,34]. More than a quarter (26%) of the farmers used antibiotics as prophylaxis as a preventive measure and 76% of farmers used them as therapeutics to treat diseases. Detection of bacterial infections in the farms even after the use of all these antibiotics is deeply concerning.

Inappropriate (and haphazard) use of antibiotics in farms can develop AMR in bacteria, making them more difficult to treat. Such usage not only leads to altered composition and diversity of gut microbiome in poultry [35,36] but also result in high levels of resistance to several classes of antibiotics, including widely used antibiotics such as Colistin, Fluoroquinolones, and Beta-Lactams [37]. Further study needs to be carried out to assess presence of AMR genes in the bacterial pathogens detected in the farms.

The World Health Organization (WHO) has developed a classification system for antibiotics called Access, Watch, and Reserve (AWaRe) to guide the appropriate use of antibiotics and combat antimicrobial resistance. The Access group contains antibiotics that should be widely available and affordable, including first-line treatments for common infections. The Watch group contains antibiotics that should be used with caution and reserved for specific indications to prevent the development of resistance. The Reserve group contains last- resort antibiotics, which should be used sparingly and only when all other options have been exhausted [38]. Apart from tetracycline, the other two antibiotics that are classified as ‘Access’ by the WHO are used by farmers sparingly (Pleuromutilin – 7% and Penicillin – 5%). Antibiotics under ‘Watch’, such as Quinolone (12%), Aminoglycoside (12%) and Macrolide (9%) are widely used by poultry farmers. Alarmingly, Colistin (19%) which is listed as a ‘Reserve’ by the WHO is the second most abundantly used antibiotics.

A complete disregard to AMR stewardship combined with lack of knowledge among poultry farmers can have devastating impact on propagation of AMR. Widespread use of strong antibiotics and indifference to implementing withdrawal period not only aggravates the AMR situation in Nepal but also pose serious food- safety risks. Withdrawal period is a withhold time prior to market distribution of poultry products after antibiotic usage in production; this is to ensure antibiotics have been degraded sufficiently and rendered inactive [39]. A survey conducted in Kathmandu of more than 200 farmers revealed only few poultry farmers knew about withdrawal periods [7] or were aware of the importance of adhering to withdrawal periods after antibiotic use to prevent the development of AMR [14,15]. All of these indicate the need of strict monitoring and control of antibiotic use by concerned government agencies.

Farmers in this study mentioned consulting either veterinarians or veterinary technicians regarding antibiotics usage (Table ***3***), however, either they did not fully comprehend the gravity of the looming AMR threat, or they disregarded the information they received. Majority of the farmers (88%) claimed to have received trusted information on AMR from various experts, but almost half of the farmers were unclear about AMR and majority (80%) of them were not willing to observe withdrawal period for their products.

In order to enhance biosecurity and reduce the risk of disease outbreaks in the poultry industry, it is necessary to raise awareness among poultry farmers about the usage of antibiotics along with significance of biosecurity measures [15,40]. This can be done through extensive practical training amongst network of poultry farmers. Proper biosecurity measures, such as the frequent disinfection of farm premises and equipment and the provision of protective clothing and footwear for workers can greatly help in prevention and containment of farm borne diseases. Additionally, reducing the use of antibiotics in poultry production and promoting alternative methods of disease prevention, such as immunization, use of probiotics and immune modulators can play a crucial role in improving poultry health and reducing disease risks.

## Supporting information

Sub Table 2

## Acknowledgement

This study was made possible by the funding support from the Regional Environment, Science, Technology and Health (ESTH) Office for South Asia- the US State Department. We would like to thank Mr. Patrick Gan, Mr. Jay Pal Shrestha, and Ms. Sulakchana Rai of ESTH office for all their support. We would also like to show our gratitude to the Department of Livestock Services. Finally, we would also like to thank laboratory associates of BIOVAC Nepal and CMDN for their help in analyzing the samples.

## Figure Captions

Figure 4: Selected poultry farms in the districts (Kavre, Sindhupalchowk, Ramechhap and Nuwakot) surrounding Kathmandu valley. The top map shows location of the districts in Nepal (shown in red border) and the bottom map shows their location in respect with Kathmandu Valley (shown just for reference, not a sampling site). The map was generated using QGIS software, Version 3.30.0

Figure 5: Graphical representation of BBCM scores received by all 16 farms for various biosafety & biosecurity parameters. Scores (percentage) received by all farms assessed in this study using BBCM score. Numbers in parenthesis in the legend refers to total number of activities assessed within each criterion.

Figure 6: Various types of antibiotics used (in %) in the surveyed poultry farms.

## Supplementary Figure Captions

Figure S 1 (A and B): Bacterial and viral diseases detected in poultry farms of Nuwakot District. The four farms were numbered from N1 to N4. Each sample represents pooled oral and cloacal samples. The gel was run with ladder in the first well and positive and negative controls in the last two well respectively. The two images show results of PCR for Mg (left, A) and IBD (right, B).

Figure S 2: IAV detected in poultry farms of Ramechhap District. The four farms were numbered from R1 to R4. Each sample represents pooled oral and cloacal samples. The gel was run with ladder in the first well and positive and negative controls in the last two well respectively.

Figure S 3: Mg detected in poultry farms of Sindhupalchowk District. The four farms were numbered from S1 to S4. Each sample represents pooled oral and cloacal samples. The gel was run with ladder in the first well and positive and negative controls in the last two well respectively.

Figure S 4 (A and B): Bacterial diseases detected in poultry farms of Ramechhap District. The four farms were numbered from R1 to R4. Each sample represents pooled oral and cloacal samples. The gel was run with ladder in the first well and positive and negative controls in the last two well respectively. The two images show results of PCR for Mg (left, A) and Ms (right, B).

Figure S 5: Mg detected in poultry farms of Kavre District. The four farms were numbered from K1 to K4. Each sample represents pooled oral and cloacal samples. The gel was run with ladder in the first well and positive and negative controls in the last two well respectively.

## Table Captions

Table 4: PCR primers for each of the pathogen designed using PrimerBlast®

Table 5: Total number and percentage of BBCM criteria met by the farms as per our assessment checklist. Overall average BBCM score was calculated by dividing the sum of all BBCM fulfilled criteria (n) and calculating the percentage. The highest BBCM score (Green cell- Sindhupalchowk Farm 4) was 67%, and the lowest BBCM score was 12% (Orange cell- Kavre Farm 3). The total number of activities assessed in each criterion are listed within parenthesis. Details of the activities assessed are shown in Supplementary Table 1. Mg – Mycoplasma gallisepticum, Ms – Mycoplasma synoviae, IBD – Infectious Bursal Disease, IAV – Influenza A Virus

Table 6: Knowledge and perception of AMR among poultry farmers. Responses (in percentage) obtained from farmers on AMR-related questions.

## Supplementary Table Captions

Supplementary Table 1: Biosafety and biosecurity checklist used to assess the farms

Supplementary Table 2: Antibiotic stewardship survey used to assess farm owners’ knowledge on antibiotics and their usage

## Notes

### Competing Interest Statement

The authors have declared no competing interest.

### Summary of Updates

We revised the acknowledgment section only.

