## Supplementary material for "Poor biosafety and biosecurity practices and haphazard antibiotics usage in poultry farms in Nepal hindering antimicrobial stewardship": Sup Table 1

**Supplementary Table 1: Biosafety and biosecurity checklist used to assess the farms**

**BIOSECURITY CHEKCLIST FOR POULTRY FARM**

**Location:** **GPS:** **Date:**

**Recorded By:**

| Activities | Completed | Remarks |
|---|---|---|
| ***General Practices*** | | |
| 1. Perimeter fence around the production area – defined biosecurity zone | | |
| 2. Any other animal or bird kept at the facility must be screened for diseases to avoid transmission | | |
| 3. Defined area for free-range poultry to graze | | |
| 4. Main entrance to the production must be capable of being closed off to other people/vehicle | | |
| 5. Defined area (away from shed) for parking, wearing PPE | | |
| 6. Entry into the shed must have a footbath or similar disinfectant available, provision for scraping boots/footwear | | |
| 7. Hand washing facility present near the shed | | |
| 8. A dedicated area for new birds and one for collection of dead birds | | |
| 9. Shed covered with mesh or tarp to prevent entry of wild birds | | |
| 10. Proper drainage system in the production area – avoid water stagnation and breeding area for diseases | | |
| 11. Baits for rodents or other pests if they frequent the farm | | |
| 12. Closed system for treated and sanitized water poultry | | |
| 13. No other birds kept in the production are apart from production birds | | |

| |
|---|
| 14. If more than one type of production bird is present, they must be housed and managed separately, shared equipment disinfected between uses |
| 15. Backyard chickens kept away from production chickens |
| 16. Closed feeding systems, protected from access & contamination by wild birds and rodents |
| 17. Any feed spilled outside the shed cleaned immediately |
| ***Personnel Standards and Procedures*** |
| 1. Poultry production area staff must NOT have any contact with other poultry, cage birds, ostriches, and pigeons while actively engaged in the production area |
| 2. Staff must wear appropriate and clean PPE or on-farm clothing and footwear |
| 3. Staff must NOT move around the farm using the same PPE, one set must be dedicated for the shed only |
| 4. Apart from staff, any other personnel must avoid multiple trips to the production area on the same day. If they must, appropriate PPE must be worn |
| 5. Hands washed/sanitized before entering the shed |
| 6. Repair/maintenance personnel or any other outside personnel must not enter populated area of the shed unless it's an emergency |
| 7. Routine maintenance and disinfection conducted between batches (for broiler chickens) |
| 8. Tools taken into production area/shed must be thoroughly cleaned and disinfected |
| 9. Records kept (logs) of all non-production staff personnel that visit the farm |

| | | |
|---|---|---|
| 10. Any other colleague, friends, or family that might have come in contact with other poultry outside of the farm must not enter the shed | | |
| 11. Non-essential vehicles must be parked at least 30m from the production area | | |
| 12. Feed transfer crews and flock pick-up crews must NOT visit on the same day | | |
| 13. Pick up vehicles must be thoroughly cleaned | | |
| 14. Day-old-chick vehicles must be disinfected before offloading the chicks and before leaving the property | | |
| 15. Delivery personnel must sanitize their hands before offloading any material | | |
| **Activities** | **Completed** | **Remarks** |
| ***Water Supply*** | | |
| 1. Water disinfected before feeding the chickens | | |
| 2. When chlorinating, the water must have minimum of 2 hours of contact time with chlorine before use | | |
| 3, Water supply must be checked for cleanliness and disinfected daily | | |
| 4. Drinking water standards - colony count $\leq$ 1000, E. Coli – Nil, Coliforms $\leq$ 100 | | |
| ***Rodent Control*** | | |
| 1. Baits placed at regular intervals, more baits placed in regions of high rodent activities | | |
| 2. Baits checked weekly and replaced with fresh baits as needed | | |
| 3. Baits designed to prevent other animals apart from rodents/pests from accessing them | | |

| |
|---|
| 4. Records kept of frequency of rodents |
| ***Cleaning and Maintenance*** |
| 1. Feed spills cleaned as soon as practicable as it attracts birds and rodents |
| 2. Grass on and around the production area are cut – long grass attracts rodents and favors survival of bacteria and viruses |
| 3. Footbaths inspected daily for excessive organic matter and replaced as required |
| 4. Free-range area adequately drained to avoid water stagnation and slope created to avoid influx of runoff water from other parts |
| 5. Manure and litter from poultry in adjacent land carefully considered to avoid spread of diseases |
| a. Dry cleaning – brushing, scraping, broom cleaning |
| b. Wet cleaning - soaking, washing, and rinsing using detergents and using washer with warm water for ceiling, walls, floors, platforms, equipment – to loosen debris and improve penetration of cleaning agents |
| c. Washing – washing every surface in the building using neutral detergent – done after wet cleaning |
| d. Disinfection – equipment disinfected within 24 hours of cleaning and drying |
| e. Disinfection – dependent on surfaces and pathogens targeted; usually work better at temperature above $20^{O}C$ |
| f. PPE worn while cleaning, washing, and disinfecting |
| ***Record Keeping*** |
| 1. Daily feed and water consumption |
| 2. Daily bird mortality or sickness |

| |
|---|
| 3. Any biosecurity breach |
| 4. Vaccinations |
| 5. Additional treatments (medications – date, duration, and outcomes) |
| 6. Visitors and vehicles logs |
| ***Others*** |
