## Supplementary material for "Poor biosafety and biosecurity practices and haphazard antibiotics usage in poultry farms in Nepal hindering antimicrobial stewardship": Sub Table 2

**Supplementary Table 2: Antibiotic stewardship survey used to assess farm owners' knowledge on antibiotics and their usage.**

**ANTIBIOTIC STEWARDSHIP SURVEY FOR FARM OWNERS**

| Questions | Remarks |
|---|---|
| Knowledge on antimicrobials? (Y/N) | |
| Prophylactic usage of antimicrobials? (Y/N) | |
| Therapeutic usage of antimicrobials? (Y/N) | |
| Conduction of post-mortem of dead animals? (Y/N) | |
| Route for antimicrobials? Feed (F), Water (W), Injection (I)? | |
| Knowledge on withdrawal period? (Y/N) | |
| Purchase with prescription? (Y/N) | |
| Withdrawal period prior to culling? (Y/N) | |
| Knowledge on Antimicrobial Resistance? (Y/N) | |
| Participation on Antimicrobial awareness programs? (Y/N) | |
| Source of antibiotic knowledge? Local animal health centre (L), Veterinary doctor(D), Veterinary technician (T), Vet suppliers and shop(S), Own(O) | |

| | |
|---|---|
| Last time antibiotic was used? This week (W), month(M), Year(Y), Never(N) | |
| Perception of AMR as a threat? Y/N | |
| More trustworthy stakeholder? Veterinary doctor(D), Vet technician (T), Local Animal Health Centre (L), Self-treatment(O), Shop and Sales (S)? | |
| Most used antibiotics according to scale | Subjective question |
